# Quantification of allosteric communications in matrix metalloprotease-1 on alpha-synuclein aggregates and substrate-dependent virtual screening

**DOI:** 10.1101/2021.01.11.426304

**Authors:** Sumaer Kamboj, Chase Harms, Derek Wright, Anthony Nash, Lokender Kumar, Judith Klein-Seetharaman, Susanta K. Sarkar

**Author notes:** These authors contributed equally (Sumaer Kamboj, Chase Harms).

## Abstract

Alpha-synuclein (aSyn) has implications in pathological protein aggregations observed in neurodegenerative disorders, including Parkinson’s and Alzheimer’s diseases. There are currently no approved prevention and cure for these diseases. In this context, matrix metalloproteases (MMPs) provide an opportunity because MMPs are broad-spectrum proteases and cleave aSyn. Previously, we showed that allosteric communications between the two domains of MMP1 on collagen fibril and fibrin depend on substrates, MMP1 activity, and ligands. However, allosteric communications in MMP1 on aSyn-induced aggregates have not been explored. Here we report quantification of allostery using single molecule measurements of MMP1 dynamics on aSyn-induced aggregates by calculating Forster Resonance Energy Transfer (FRET) between two dyes attached to the catalytic and hemopexin domains of MMP1. The two domains of MMP1 prefer open conformations, with the two domains well-separated. These open conformations are inhibited by a single point mutation E219Q of MMP1 and tetracycline, an MMP inhibitor. A two-state Poisson process describes the interdomain dynamics. The best-fit parameters for a Gaussian fit to the distributions of FRET values provide the two states. The ratio of the kinetic rates between the two states comes from the ratio of fitted areas around the two states. The decay rate of an exponential fit to the correlations between FRET values provides the sum of the kinetic rates. Since a crystal structure of aSyn-bound MMP1 is not available, we performed molecular docking of MMP1 with aSyn using ClusPro. We simulated MMP1 dynamics using different docking poses and matched the experimental and simulated interdomain dynamics to determine the most appropriate pose. We performed virtual screening against the potential ligand-binding sites on the appropriate aSyn-MMP1 binding pose and showed that lead molecules differ between free MMP1 and substrate-bound MMP1. In other words, virtual screening needs to take substrates into account for substrate-specific control of MMP1 activity. Molecular understanding of interactions between MMP1 and aSyn-induced aggregates may open up the possibility of degrading pathological aggregates in neurodegeneration by targeting MMPs.

**Significance:** We have quantified MMP1 interdomain dynamics on aSyn-induced aggregates by a two-state Poisson process. Histograms and correlations of FRET values determine the kinetic rates of interconversion between the two states. We quantify the conformational dynamics of the whole MMP1 and allosteric communications by the two-dimensional matrix of correlations between every pair of amino acids from experimentally-validated all-atom simulations. The two-dimensional correlations lead to a Gray Level Co-occurrence Matrix and a measure of Shannon entropy describing the conformational fluctuations. As such, we address the quantification of allosteric communications, a leading challenge in defining allostery. We report that the potential ligand-binding sites and lead molecules change for MMP1 upon binding alpha-synuclein and depend on the binding pose selected. This suggests that one needs to take the substrate into account while targeting MMPs.

## Introduction

Lewy bodies, intraneuronal cytoplasmic protein aggregates, are commonly observed in the brains of patients with Alzheimer’s disease (AD) and Parkinson’s disease (PD) [1]. Alpha-synuclein (aSyn) is often the common protein detected in Lewy bodies observed in AD and PD [2]. An established body of evidence shows that the C-terminal truncation of aSyn by proteases leads to protein aggregation and Lewy body formation. Both intracellular and extracellular proteolytic systems take part in the breakup of aSyn [3]. As such, partial cleavage of aSyn explains aggregates due to fibrillation [4] and misfolding [5, 6]. Although aggregates could be a consequence instead of a cause of AD and PD [7, 8], the uncontrolled aggregation will lead to cell death and worsening of effects [9, 10]. As such, there is a need to find ways to degrade existing aggregates.

A growing body of evidence shows that matrix metalloproteases (MMPs) can partially cleave aSyn [11–15]. Prior reports have shown that MMP1, MMP2, MMP3, MMP9, and MT1-MMP can partially cleave aSyn [13, 14] with the possible cleavage sites identified [13]. MMPs are broadspectrum proteases known to degrade extracellular matrix and non-matrix proteins [16, 17], but a growing body of evidence suggests proteolytic and non-proteolytic intracellular functions of MMPs [18]. MMPs are found in extracellular space [19], intracellular space [20], blood [21], intestine [22], brain [23], and can alter the blood-brain barrier [24]. The implication of MMPs in partial cleavage of aSyn is significant because tetracycline [25, 26], a well-known inhibitor of MMPs [27], have shown therapeutic potential in AD [28] and PD [29]. The broad-spectrum protease activity of MMPs not only leads to aSyn-induced aggregation, but it might also be useful in degrading existing aSyn-induced aggregates. However, it is not clear yet how MMPs might affect aSyn-induced aggregates, which are water-insoluble and challenging to study using standard biochemical assays. To this end, we have developed a single molecule tracking approach and weight-based activity assay to study water-soluble MMPs interacting with water-insoluble substrates. As such, it is possible to study MMP activity on aSyn-induced aggregates at the single molecule level.

However, it is not enough to know how MMP interacts with aggregates. Due to broad-spectrum activity, targeting MMPs for improving human health is challenging because MMPs interact with and degrade many proteins in the human body [30]. Due to such diverse functions, any drug used for inhibiting MMPs results in adverse side effects [31]. As such, there is only one FDA-approved MMP inhibitor (doxycycline hyclate) [32]. To circumvent this problem, we need to understand the mechanisms behind diverse functions. The catalytic domain sequence of MMP1 is very similar to other MMPs in the family. However, the hemopexin domain sequence varies [33, 34], suggesting that differences in activity and substrate specificity among MMPs are due to allosteric (distant from the catalytic site) communications [17, 35]. Recently, we reported activity-dependent MMP1 dynamics and allosteric communications on type-1 collagen fibrils [36] and fibrin [37]. We showed that functionally-relevant conformations have the catalytic and hemopexin domains of MMP1 well-separated. These open conformations often accompany a larger catalytic pocket opening of MMP1, enabling the substrates to get closer to the active site of MMP1. Building on this, we have the opportunity to define dynamics and allosteric communications of MMP1 on aSyn-induced aggregates.

In this paper, we report measurements of activity-dependent MMP1 interdomain dynamics on aSyn-induced aggregates of *E. coli* proteins at the single molecule level. MMP1 prefers open conformations on aSyn-induced aggregates, and these conformations are inhibited by tetracycline. A two-state Poisson process describes the interdomain dynamics of MMP1 on aggregates. Since aSyn-bound MMP1 crystal structures are not available, we used ClusPro for molecular docking of MMP1 with aSyn. We performed all-atom simulations of different binding poses to calculate the interdomain dynamics and compare them with experimental dynamics. It appears that the pose with MMP1 bound to the middle U-shaped region of aSyn structure leads to dynamics similar to experimental dynamics. We used this experimentally-guided binding pose to quantify allosteric communications. We also determined the potential ligand binding sites and performed virtual screening against one of these binding sites to compare the lead molecules for free and aSyn-bound MMP1, which showed substrate-dependence of virtual screening. Our results pave the way for substrate-dependent virtual screening guided by molecular insights that may enable inhibition of aggregation and degradation of aggregates.

## Results and discussion

### MMP1 dynamics on aSyn-induced aggregates show activity-dependent conformations and correlations

We purified catalytically active (E219) and inactive (Q219) MMP1 using a protease-based method described in our previous publication. Both active and inactive MMP1 had SER142CYS mutation in the catalytic domain and SER366CYS mutation in the hemopexin domain for labeling with the Alexa555 (donor)-Alexa647 (acceptor) FRET pair. Labeling does not affect the activity of MMP1. To distinguish the effects of enzymatic activity despite potential problems due to the labeling dyes’ photophysical properties, we used inactive MMP1 as control. Since aSyn-induced aggregates are water-insoluble, we created a thin layer of aggregates on a quartz slide. We flowed water-soluble MMPs to image the dynamics of MMP1 using a TIRF microscope. We used a laser at 532 nm to excite Alexa555 (see methods). MMP1 undergoes interdomain dynamics and the distance between Alexa555 and Alexa647 changes. As a result, the non-radiative energy transfer due to FRET between the two dyes changes. Low FRET conformations lead to high Alexa555 emission, whereas high FRET conformations lead to low Alexa555 emission. Anticorrelated Alexa647 and Alexa555 emissions, I_A_ and I_D_, respectively, indicate the conformational dynamics of MMP1. We calculated FRET using the equation IA/(IA+ID), where each FRET value determines the separation between the two MMP1 domains. We plotted the distribution of FRET values to determine the MMP1 conformations. We calculated the correlation between conformations to determine how a conformation at one time is related to another conformation at a later time. A sum of two Gaussians fits the histograms, whereas an exponential fits the correlations. We fitted both power law and exponential to the autocorrelations because these are the most common decay types of correlations.

### A two-state Poisson process describes MMP1 dynamics on aSyn-induced aggregates

Recently, we published a quantitative analysis of MMP1 dynamics on collagen fibrils [36] and fibrin [37]. The histograms reveal conformations of MMP1, and the autocorrelations show the relation between conformations at different time points. A sum of two Gaussians fits the histograms of smFRET values, and an exponential fits the autocorrelations. As such, a two-state Poisson process describes the conformational dynamics of MMP1 and enables a straightforward interpretation of the decay rates of correlations [36]. We can establish a quantitative connection between the two states obtained from the Gaussian fits and the exponential fits’ decay rates. We defined the two Gaussian centers as the two states, S1 (low FRET) and S2 (high FRET). Also, we defined the two kinetic rates as k1 (S1→S2) and k2 (S2→S1) for interconversion between S1 and S2. For a two-state system, the ratio of rates (k1/k2) is the ratio of Gaussian area(S2)/area(S1), and the sum of rates (k1+k2) is the decay rate of autocorrelation. We calculated k1 and k2 from k1/k2 and k1+k2 for both active and inactive MMP1. Using experimentally-determined S1, S2, k1, k2, and noise (widths of the histograms), we simulated smFRET trajectories and analyzed the simulated data similar to the experimental data. Comparing experimentally-determined inputs (**Table S1**) and recovered parameters from two-state simulations (**Table S2**) shows that a two-state Poisson process describes MMP1 interdomain dynamics.

The two states are S1=0.46 and S2=0.52 on aSyn-induced aggregates for active MMP1 without ligand (**Table S1**). In comparison, the two states are S1=0.44 and S2=0.55 on collagen [36] and S1=0.42 and S2=0.51 on fibrin [37] for active MMP1 without ligand. The correlation decay rate of 0.08 s^-1^ on aSyn-induced aggregates for active MMP1 without ligand (**Table S1**). In comparison, the decay rates are 0.13 s^-1^ on collagen [36] and 0.08 s^-1^ on fibrin [37] for active MMP1 without ligand. In the presence of tetracycline, the two-state description still applies even though the two states and interconversion rates between them change compared to the dynamics without ligand (**Table S1**). We performed two-state simulations with and without noise to study how noise affects the histograms and autocorrelations. Both power law and exponential fit the autocorrelations of simulated smFRET trajectories without noise (**Figure 2F**). However, only an exponential fits the autocorrelations with noise. In other words, the presence of noise can convert a power law correlation into an exponential correlation. This effect is similar to the conversion of a Lorentzian line shape into a Gaussian line shape [38]. Note that an exponential fit recovers the simulated kinetic rates’ underlying sum with and without noise (**Table S2**).

**Figure 1.**
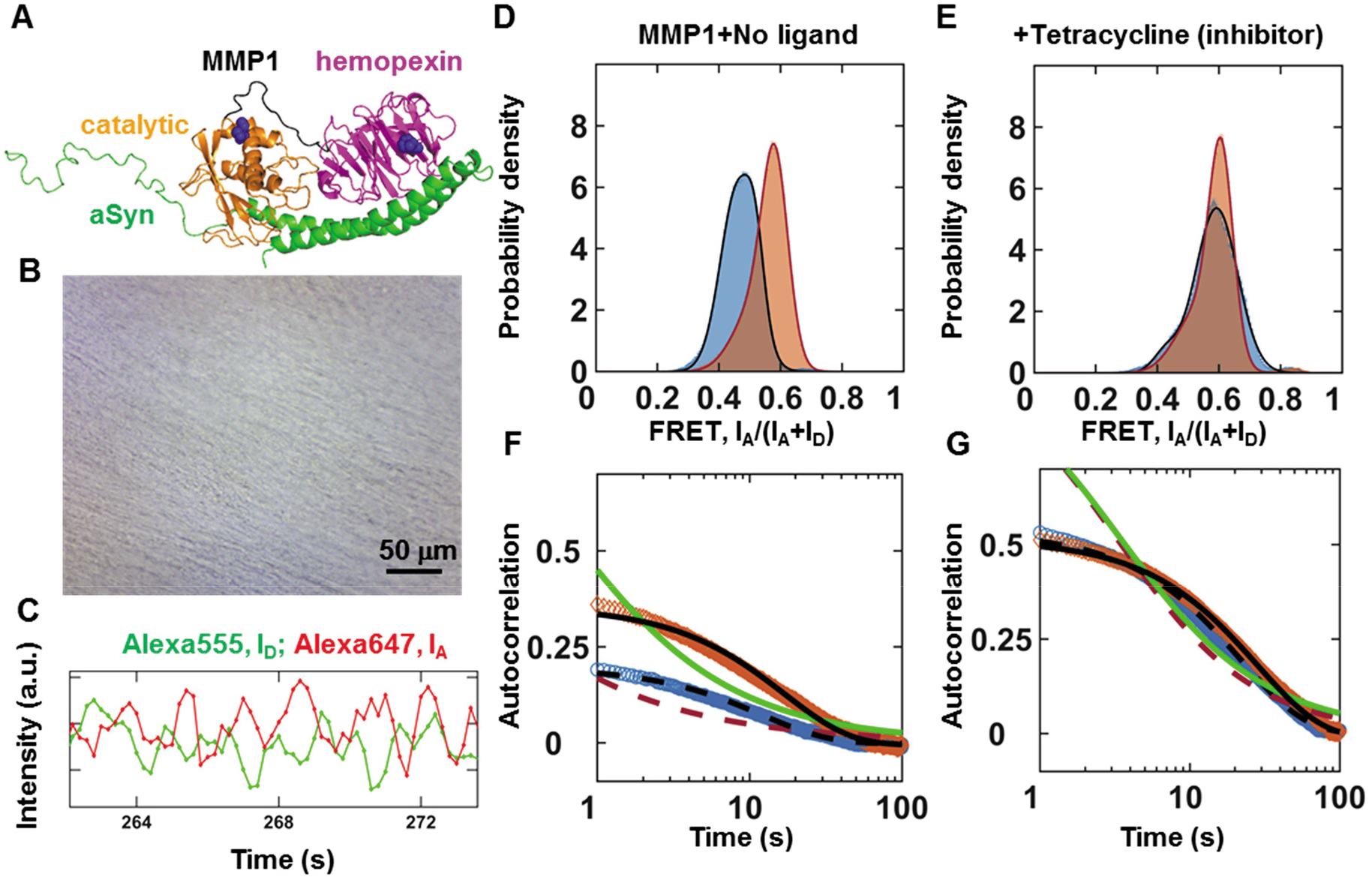
Activity-dependent interdomain dynamics of MMP1 on aSyn-induced aggregates at 22 °C with 100 ms time resolution. (**A**) Relative positions of the MMP1 domains and residues created using PDB ID 4AUO. (**B**) Light microscopy image of aSyn-induced aggregates on a slide. For MMP1-treated aggregates, see **Figure S1**. (**C**) Emission intensities of the two dyes attached to active MMP1. (**D**) and (**E**) Area-normalized histograms of MMP1 interdomain distance (bin size=0.005) without ligand and in the presence of tetracycline (an inhibitor), respectively, for active (blue) and inactive (orange) MMP1. All histograms are fitted to a sum of two Gaussians (active: solid blue line; inactive: solid red line). (**F**) and (**G**) Autocorrelations of MMP1 interdomain distance without ligand and in the presence of tetracycline, respectively, for active (blue) and inactive (orange) MMP1. All autocorrelations are fitted to exponentials and power laws (exponential fit to active: dashed black line; power law fit to active: dashed red line; exponential fit to inactive: solid black line; power law fit to inactive: solid green line). The error bars in histograms and autocorrelations represent the square roots of the bin counts and the standard errors of the mean (SEM) and are too small to be seen. For best-fit parameters for histograms and autocorrelations, see **Table S1**.

**Figure 2.**
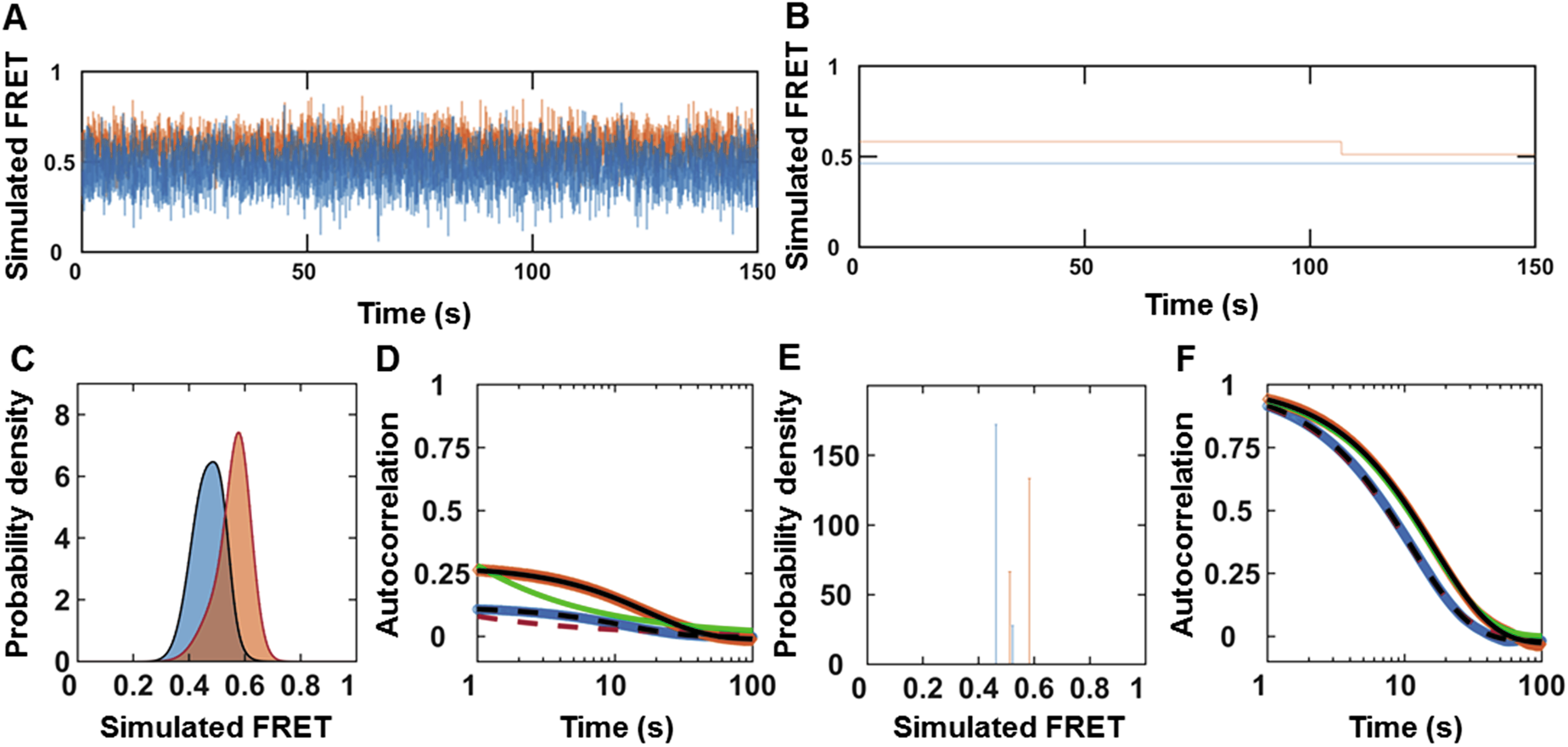
Stochastic simulation of MMP1 interdomain dynamics as a two-state system. (**A**) and (**B**) Examples of simulated smFRET trajectories with and without noise, respectively, for active MMP1 (blue) and inactive MMP1 (orange) using experimentally-determined parameters for MMP1 without ligands. (**C**) Area-normalized histograms of simulated smFRET values with noise (active: blue; inactive: orange) with best fits to a sum of two Gaussians (solid black line). (**D**) Autocorrelations of simulated smFRET trajectories with noise (active: blue; inactive: orange) with best fits to exponentials (active: dashed black line; inactive: solid black line). Power law does not fit autocorrelations (active: dashed red line; inactive: solid green line). (**E**) Area-normalized histograms of simulated smFRET values without noise (active: blue; inactive: orange) with best fits to a sum of two Gaussians (solid black line). (**F**) Autocorrelations of simulated smFRET trajectories without noise (active: blue; inactive: orange) with best fits to exponentials (active: dashed black line; inactive: solid black line). Both exponential and power law fit autocorrelations (active: dashed red line; inactive: solid green line). The error bars are the SEMs for histograms and autocorrelations and are too small to be seen. For best-fit parameters, see **Table S2**.

### MMP1 dynamics depend on the aSyn-MMP1 binding pose

Since crystal structures of aSyn-bound MMP1 do not exist, we determined the binding poses of aSyn (PDB ID 1XQ8) and MMP1 (PDB ID 4AUO) computationally using molecular docking software ClusPro [39, 40]. **Figures 3A-C** show three such binding poses. We selected these poses based on known cleavage sites of aSyn targeted by MMP1 because the computational binding energy may not be the best indicator of appropriate docking poses [41]. We performed all-atom MD simulations for the three binding poses (**Figure 3**) and calculated the catalytic pocket opening (**Figures 3D-F**) and interdomain separation (**Figures 3G-I**) of MMP1. Since MMP1 stabilized within ~5 ns (**Figure S2**), we performed simulations for 20 ns. Previously, we reported a positive correlation between the catalytic pocket opening and interdomain separation for MMP1 dynamics on collagen [36] and fibrin [37]. Based on the criterion of positive correlation, it appears that binding pose 3 is likely to be one of the most appropriate binding poses. Previous reports of MMP1 cleavage sites on aSyn is consistent with our suggestion that pose 3 is a suitable binding pose.

**Figure 3.**
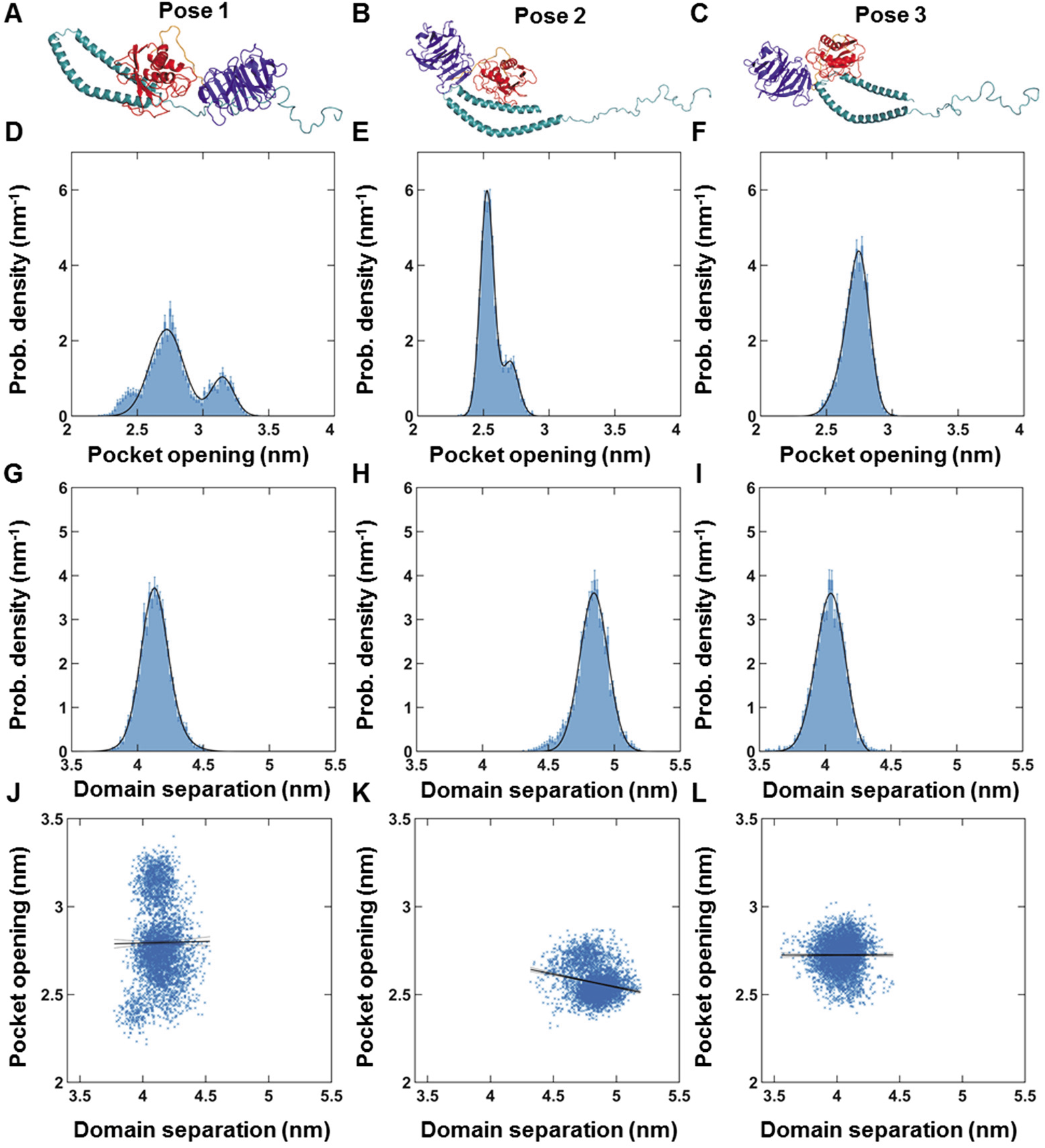
MMP1-aSyn binding pose-dependent dynamics of active MMP1 at 37 °C. (**A**), (**B**), and (**C**) Three binding poses of MMP1 at different locations of aSyn. (**D**), (**E**), and (**F**) Area-normalized histograms of the catalytic pocket opening of MMP1 for poses 1, 2, and 3, respectively. (**G**), (**H**), and (**I**) Area-normalized histograms of interdomain separation of MMP1 for poses 1, 2, and 3, respectively. (**J**), (**K**), and (**L**) Linear correlation plots of catalytic pocket opening and interdomain distance for poses 1, 2, and 3, respectively. The data were fitted to *y_i_* = *b*_0_+*b*_1_×*x_i_*. Note that a larger domain separation corresponds to a lower FRET value. Time resolution=2 fs, Data saved every 5 ps, RMSD stabilization time for MMP1=~5 ns, Total simulation duration=20 ns. For calculations of linear correlations, see methods. For best-fit parameters, see **Table S3**.

### Shannon entropy enables quantification of allosteric communications

Proteins are inherently flexible biomolecules having both intra- and interdomain motions. Two types of randomness appear as proteins interact with their surroundings and substrates. At a particular time point, the amino acids’ locations and angles have a spatially random component. As time progresses, proteins sample through different conformations leading to temporal randomness. Such correlated or collective fluctuations are essential for functions, including allosteric regulation [42–44], generation of mechanical work [45, 46], catalysis [47], ligand binding [48], and protein folding [49]. Correlated allosteric motions decrease the conformational entropy and affect the kinetics and thermodynamics of biological processes [50]. Shannon entropy is a quantitative measure to describe randomness in computer science and pattern recognition and has been successfully applied to quantify allostery in proteins to gain insights into their functional relevance [51–54].

Since a correlated motion suggests a decrease in randomness or lower entropy, we calculated correlations between each pair of residues in MMP1 and estimated entropy (**Figure 4**). We divided all-atom simulations into 1 ns long windows. In each 1 ns window, there were 200 radial coordinates for residues. We calculated the correlations between every pair of residues and normalized to a range of 0 to 1 by subtracting the minimum and then dividing by the maximum. **Figures 4A-C** show the matrix of correlation values at lag number 1, averaged over 20 ns. We divided the correlation values to create 10 bins of width 0.1, calculated 10×10 gray-level cooccurrence matrix (GLCM), and defined Shannon entropy*S* = –∑ *p_i_* ln *p_i_*, where *p_i_* is the probability of a microstate *i*. The catalytic domain residues (F100-Y260) have strong correlations with the hemopexin domain residues (D279-C466), suggesting allosteric communications in MMP1 (**Figures 4A-C**). The time-evolutions of Shannon entropy are shown in **Figures 4D-F**. Pose 3 has the lowest entropy consistent with the functionally relevant binding pose.

**Figure 4.**
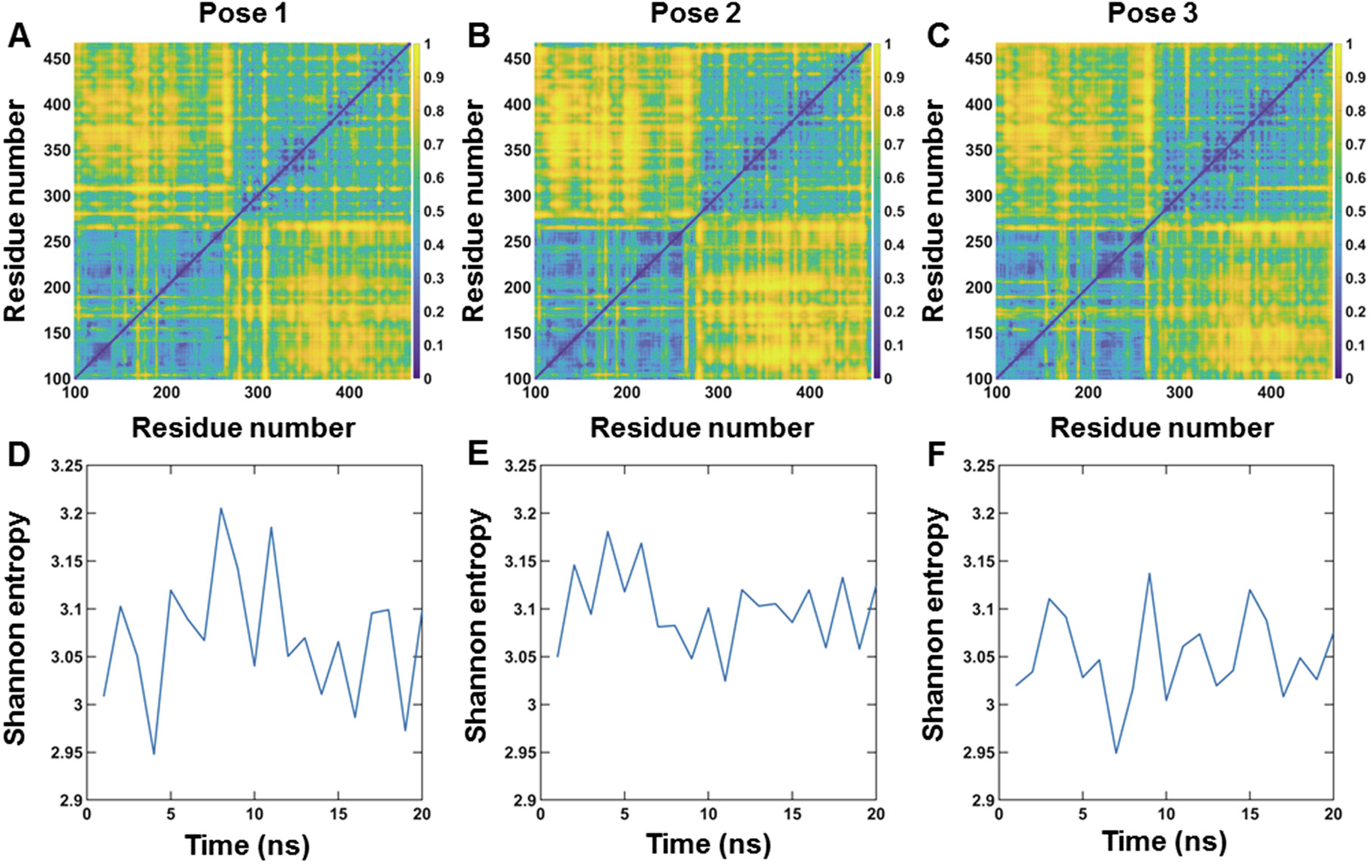
Pose-dependent correlation between residues of active MMP1 and quantification of allostery using Shannon entropy. (**A**), (**B**), and (**C**) Correlations between residues for poses 1, 2, and 3, respectively, at 37 °C. (**D**), (**E**), and (**F**) Shannon entropy calculated from correlation plots for poses 1 (S=3.07±0.02, mean±SEM), 2 (S=3.10±0.01, mean±SEM), and 3 (S=3.05±0.01, mean±SEM), respectively. For details, see supplementary information and **Figure S3**.

### Experimentally-measured dynamics agree with MD simulation dynamics

We performed allatom MD simulations of the binding pose 3 at 22 °C for active and inactive MMP1. **Figure 5A** shows the distributions of the interdomain distance between S142 and S366. In agreement with experiments (**Figure 1D**), active MMP1 adopts conformations with the two domains separated more than inactive MMP1. Also, simulated dynamics correlations (**Figure 5B**) follow the experimental pattern (**Figure 1F**), with inactive MMP1 having higher values with longer correlation times. We used experimentally-consistent simulations to gain further insights. First, the correlation between the catalytic pocket opening and interdomain separation for active MMP1 is higher than inactive MMP1 (**Figure 5C**). A larger catalytic pocket opening enables substrates to get closer to the active site. Second, the two-dimensional correlation plots show allosteric communications between the two domains (**Figures 5C-D**). Third, Shannon entropy describing the randomness of two-dimensional correlations plots shows a lower value for active MMP1 than inactive MMP1 (**Figure 5F**).

**Figure 5.**
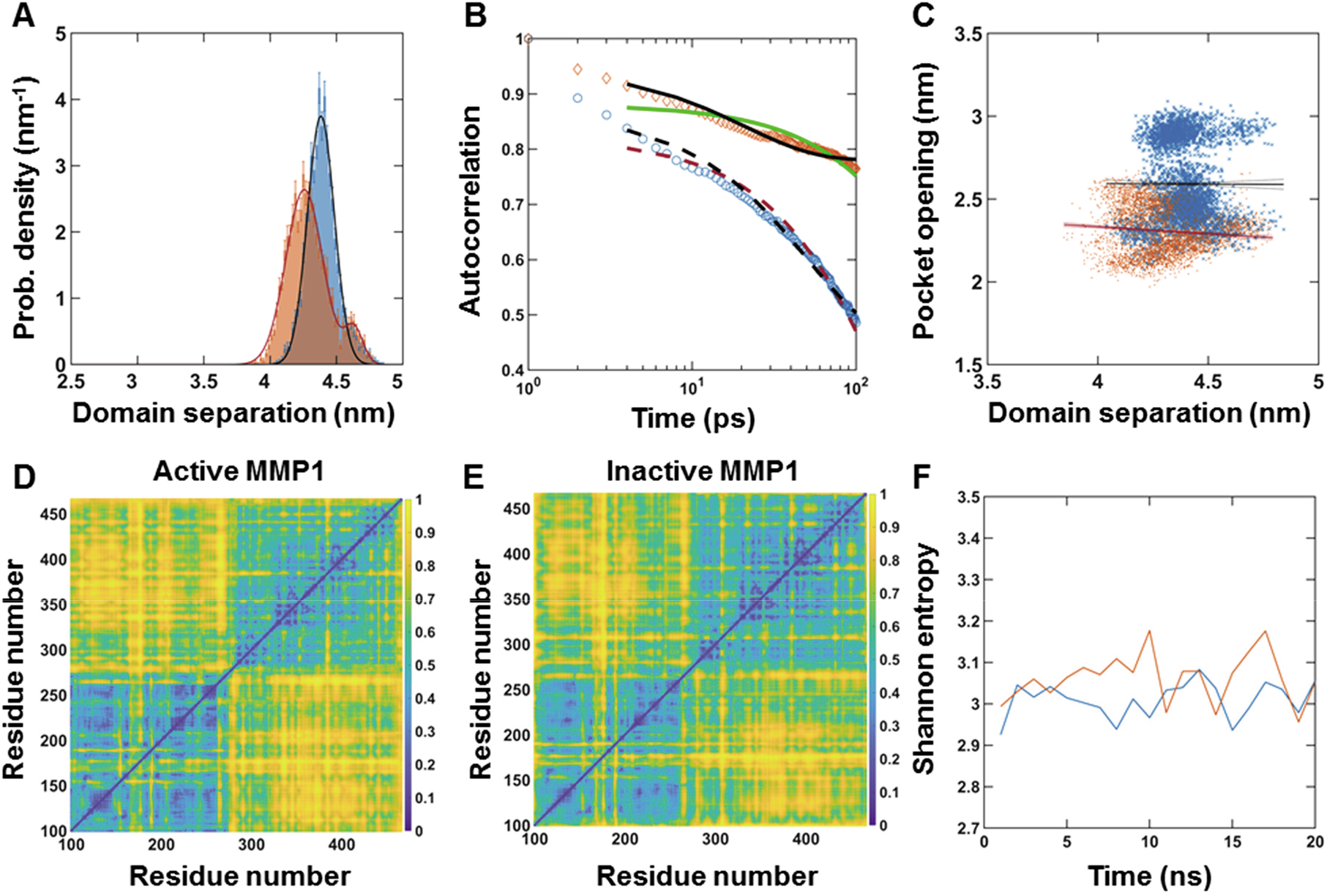
All-atom MD simulation of MMP1 interdomain dynamics at 22 °C. (**A**) Area-normalized histograms of simulated interdomain distance (active: blue; inactive: orange) with best fits to a sum of two Gaussians (solid black line). (**B**) Autocorrelations of simulated interdomain distance (active: blue; inactive: orange) with best fits to exponentials (active: dashed black line; inactive: solid black line). Power law does not fit autocorrelations (active: dashed red line; inactive: solid green line). (**C**) Linear correlation plots of catalytic pocket opening and interdomain distance. (**D**) and (**E**) Correlations between residues for active and inactive MMP1, respectively. (**F**) Shannon entropy calculated from correlation plots for active (S=3.01±0.01, mean±SEM) and inactive (S=3.06±0.01, mean±SEM). For best-fit parameters, see **Table S4**.

### Lead molecules from virtual screening against MMP1 differ upon substrate binding

Guided by experiments and simulations, we determined that pose 3 is likely a relevant binding pose between MMP1 and aSyn. We used the experimentally-informed binding pose for the virtual screening of molecules. Virtual screening enables testing considerably more ligands economically and faster than high-throughput experimental screening. A wide range of strategies can be divided into ligand-based and structure-based virtual screening [55]. In ligand-based virtual screening, known ligands against a target serve as benchmarks to screen for more ligands with similar properties. In contrast, structure-based screening uses the binding sites on a target to screen molecules leveraging the conformational changes and energetic complementarity upon ligand binding. Since MMP1 has broad-spectrum protease activity, we determined the binding sites on free MMP1 and aSyn-bound MMP1 to investigate a substrate’s effects. We predicted the potential ligand-binding sites using AutoLigand and MGLTools 1.5.7 and selected a potential binding site in the hemopexin domain of MMP1 to screen for allosteric ligands (**Figure 6A**).

**Figure 6.**
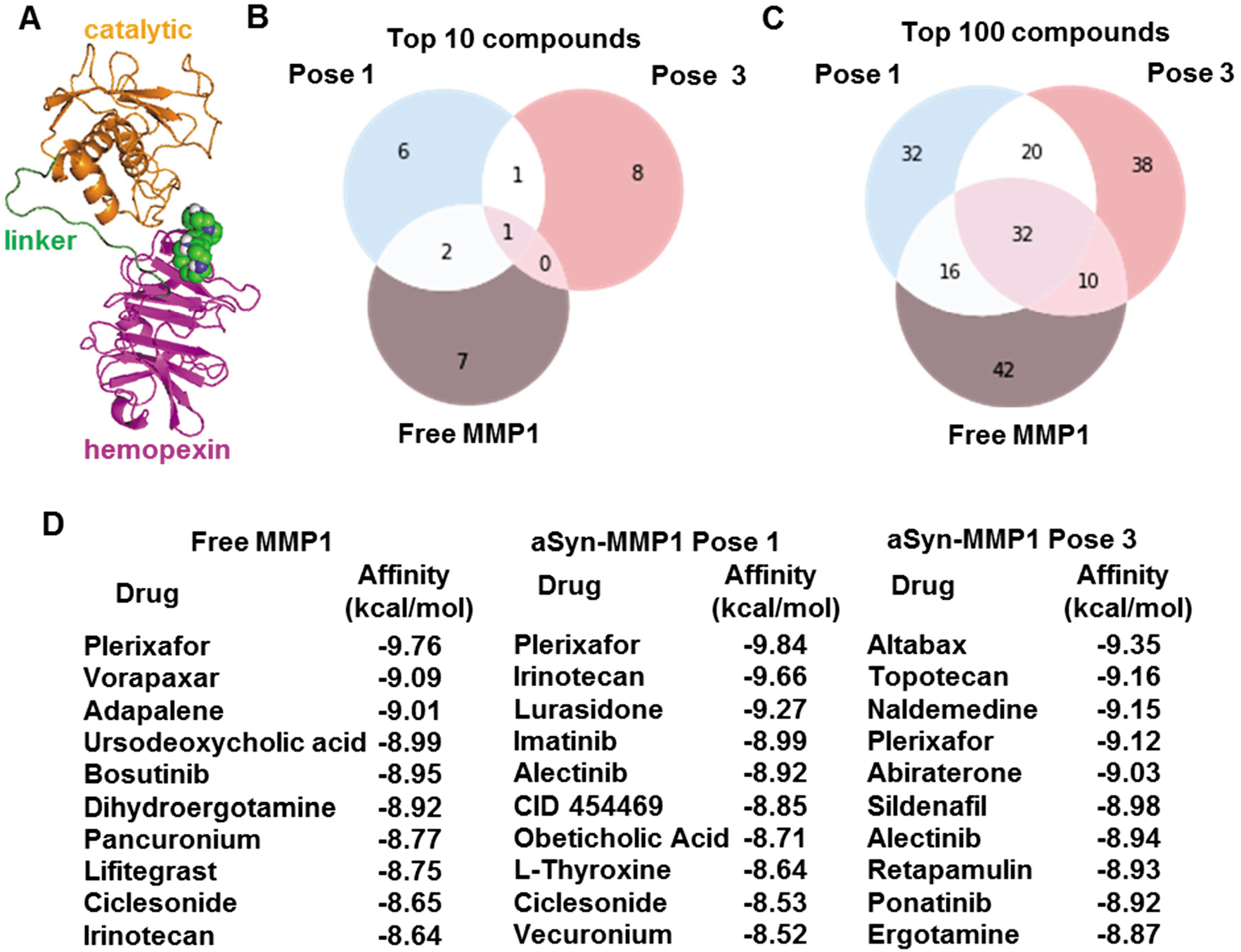
Substrate- and pose-dependent virtual screening against MMP1. (**A**) MMP1 bound to Plerixafor (green and blue spheres), the top hit against free MMP1. We performed virtual screening against the same site for free MMP1, pose 1, and pose 3. (**B**) and (**C**) Venn diagrams of top 10 and 100 molecules show unique and common ligands. (**D**) Identities and affinities for the top 10 molecules. For docking parameters, see supplementary information.

We performed screening against the selected site in the hemopexin domain for free MMP1 and two poses of aSyn-bound MMP1. We performed molecular docking using AutoDock and set up the virtual screening with Raccoon 1.1. We used default AutoDock docking parameters (see supplementary information) to determine the binding affinities for a collection of 1,400 FDA-approved compounds from the ZINC15 database. We used FDA-approved molecules because those provide a smaller set of commercially available molecules. We fixed the side chains that are otherwise flexible. Flexible side chains would have given slightly more accurate screening but would require more computational time. We found that the lead molecules against MMP1 change depending on the substrate and pose. **Figures 6B-D** show the top molecules for free MMP1 and the two aSyn-MMP1 binding poses, suggesting that substrate binding and different poses can lead to different results. Venn diagrams of molecules (**Figure 6A**) show that there are molecules unique to each case. The ligand-binding site was common to free MMP1, pose 1, and pose 3. However, conformational differences in the three cases clearly impact virtual screening. The unique molecules for each case suggest that we should perform substrate-specific screening to identify unique ligands, thus resolving problems arising from MMP promiscuity.

In summary, we measured the interdomain dynamics of MMP1 on aSyn-induced protein aggregates and modeled the dynamics as a two-state Poisson process. Distributions of conformations and correlation decay rates of MMP1 on aSyn-induced aggregates follow the general pattern that we previously reported for collagen [36] and fibrin [37], i.e., open MMP1 conformations are functionally relevant, and there are time-dependent correlations of conformations.

Since there is no crystal structure for aSyn-bound MMP1, we determined the binding poses using molecular docking. We performed all-atom simulations of dynamics for different binding poses between aSyn and MMP1 and compared them with single molecule measurements of dynamics. A comparison of experiments and simulations suggest that the pose in which MMP1 binds to aSyn near TYR39 (pose 3) leads to a better match with experiments. However, there are two important caveats. First, we performed simulations on aSyn monomer, but the experiments were done on aggregates. We took a similar approach for collagen and found that simulations on collagen monomer agreed with experiments on collagen fibril when the collagen backbone was restrained [36], suggesting strains in collagen monomers inside collagen fibrils. For aSyn, we did not have to restrain the aSyn backbone for agreement between simulations and experiments, suggesting a lack of strain in aSyn-induced aggregates. Second, aSyn is considered an intrinsically disordered protein [56] that assumes structure upon binding substrate [57]. Also, aSyn purified from neuronal and non-neuronal cell lines generally suggests a “natively folded” ~58 kDa tetrameric form [58, 59]. Nevertheless, the combined experiment-simulation approach using the monomer structure of aSyn provides a starting point for molecular-level understanding.

We used experimentally-guided simulations to quantify the randomness of dynamics by calculating correlations between each pair of residues in MMP1. We defined another matrix called GLCM to define Shannon entropy from the correlation matrix to quantify the randomness with a single number at each time point. The entropy for active MMP1 is smaller than inactive MMP1, suggesting that MMP1 activity may be entropy-driven because a lower entropy (stabilized conformations) is likely essential for substrates to get closer to the active site for a long enough duration for catalysis.

We performed virtual screening of molecules against aSyn-bound MMP1 and compared it with free MMP1. To this end, we found that the potential ligand-binding sites change upon binding aSyn and depend on the binding pose. As such, we selected an allosteric binding site on MMP1 and screened 1,400 FDA-approved molecules against the site for free MMP1 and two binding poses with aSyn. The lead molecules and their scores change upon binding the substrate and depend on the binding pose. Therefore, substrate-dependent binding sites and lead molecules suggest that any effort to target MMP1 may need to consider the substrate, the binding pose, and the ligand-binding site. The synergistic combination of experiments and simulations enables modulation of MMP1 activity using molecular insights at the single molecule level. It may pave the way for substrate-specific control of MMP activity using allosteric ligands.

## Materials and methods

### Purification of MMP1 and aSyn

The MMP1 and aSyn sequences were inserted into the pET21b+ and pET11a vectors, respectively, between NdeI (N-terminal) and HindIII (C-terminal) restriction sites. We transformed the plasmids into Rosetta (DE3) pLysS *E. coli* (Millipore, Cat# 70956-4) for protein expression. We purified MMP1 and aSyn, as described in our previous publications [60, 61]. The method of purifying aSyn also produced aSyn-induced aggregates used in experiments.

### Measurements of MMP1 interdomain dynamics

For smFRET measurements of MMP1 interdomain dynamics, we mutated two serine residues in MMP1 at the locations 142 and 366 to cysteines for labeling with Alexa555 and Alexa647 dyes. In addition, we introduced the E219Q mutation in the catalytic domain to create a catalytically inactive mutant of MMP1. We spread aSyn-induced aggregates and made a thin layer on a quartz slide. We made a flow cell for single molecule experiments using a piece of double-sided adhesive tape sandwiched between the quartz slide and a glass coverslip. Labeled MMPs were flowed into the flow cell and excited at 532 nm wavelength using the evanescent wave created at the quartz slide and sample buffer interface in a Total Internal Reflection Fluorescence (TIRF) microscope. We acquired two-channel movies to detect emissions from Alexa555 and Alexa647. Any relative motion between the two MMP1 domains would lead to a non-radiative transfer of energy from Alexa555 to Alexa647 due to FRET, increasing the emission from Alexa647 (IA) and simultaneously reducing the emission from Alexa555 (ID). We calculated FRET efficiency by IA/(IA+ ID) [62]. Single-molecule experiments and analyses have been described in our previous publications [63–66].

### All-atom simulations

We removed all zinc and calcium ions and the water molecules from PDB ID 4AUO and replaced the missing side-chain atoms using Chimera’s rotamer tool. PDB files for active and inactive MMP1 were created by replacing A219 with E219 and Q219, respectively, using Pymol’s mutagenesis function. We used Gromacs 2019.6 with the Gromos96 43a1 force field to perform the MD simulations. We ignore the hydrogen atoms while creating the topology file. Each simulation ran 20 ns long, simulated at 2 fs/step, and sampled every 5 ps. Each MMP1 crystal was placed in a cubic box with 3D period boundary conditions and solvated with water (using a single point charge model (SPC)) and Na counter-ions to create a neutral system. We then used the steepest descent algorithm to minimize the solution’s energy. To equilibrate the solution with the protein complex, we used NVT and NPT ensemble simulations. First, we performed the NVT simulation and set the mean temperature at the desired temperature of 295 K (22 °C) or 310 K (37 ° C) using a Berendsen thermostat for 100 ps. We used the Verlet cut-off scheme for neighbor searching and updated the neighbor list every 20 fs. We used the particle mesh Ewald scheme to calculate the electrostatic interactions with a cubic interpolation order of 4 and a cut-off at 1 nm. We assigned initial velocities using a Maxwell distribution from the corresponding temperature. In the NPT simulation, we maintained velocities from the NVT simulation output, and the pressure was set to 1 bar using a Parrinello-Rahman barostat for an additional 100 ps. Once the NVT and NPT simulations finished and the system equilibrated, we removed the position restraints and ran the production MD for 20 ns. We edited the final coordinates in the trajectory file to correct for periodicity and center the protein complex. The distances between the catalytic pocket residues and the serine residues (interdomain distance) are then measured at every time step using the Gromacs distance function. We measured the interdomain separation between the nitrogen atoms of residues S142 and S366 and the catalytic pocket opening between the nitrogen atoms of residues N171 and T230. To calculate the correlation and Shannon entropy, we recorded the nitrogen atom’s coordinates in each residue of MMP1.

### Analysis of experimental and simulated interdomain dynamics

We chose the histograms’ bin width to be at least the inverse of the sample size’s square root, i.e., 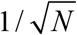, N is the number of data points. We calculated the error of count in each bin as the square root of the bin count. Both the bin counts and errors were divided by the area of the histogram to create the area-normalized histogram (probability density function). The area under the normalized histogram equals 1. We fitted a sum of two Gaussians to the histograms:

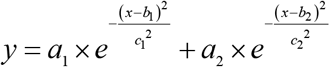

where a’s, b’s, and c’s are amplitudes, centers, and widths of the Gaussians. The parameters b1 and b2 are the two states, S1 and S2.

To calculate correlations, we subtracted the average value from each trajectory and used the following equation to calculate correlations:

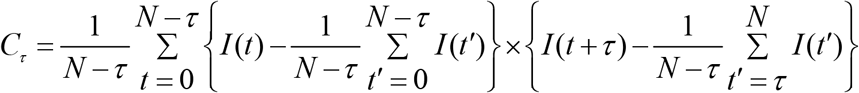

where *C_τ_* is the correlation at lag number *τ, N* is the number of points in a FRET trajectory, and *I*(*t*) is the FRET value at *t*. For autocorrelations, both factors in curly brackets were from the same time series. For cross correlations, the two factors in curly brackets were from different time series.

We normalized correlations by dividing correlation values at each lag by *C*_*τ*=0_. We fitted correlations between *τ* = 1 and *τ* = 1000 to both power law and exponential distributions. For power law, we used a form of Pareto distribution **[67]** that satisfies the boundary conditions, i.e., *C*_*τ* = 0_ = 1at *t* = 0 and *C*_*τ* = ∞_ = 0 at *t* = ∞.

We fitted the following equations of power law and exponential functions:

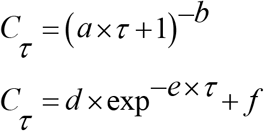

To quantify correlations between the catalytic pocket opening and interdomain separation from simulations, we used the interdomain distance as the single predictor variable for the catalytic pocket distance in a linear model. A linear model describes a response variable as a function of predictor variables. Linear models are often fit using a method known as a statistical regression. There are different regression methods used, depending on the number of predictor variables. When using only one predictor variable, the regression method is known as the simple linear regression, which we used for quantifying correlations. We used the following equation:

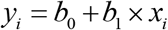

where *b_0_* and *b_1_* are the estimated fit parameters. There are also a few different methods to estimate the parameters of the linear model. The most-used approach is the least-squares operator, which finds the slope through the data that minimizes the squared distance between the fit and the residues. There is still uncertainty in these estimations, no matter which method we used. Confidence bands visually represent the uncertainty in linear models by showing the range of possible slopes. We calculated the 95% confidence interval of each predictor value’s mean.

### Small molecule putative binding sites on MMP1

For virtual screening, we selected free MMP1 and two poses of aSyn-MMP1 binding. First, we predicted potential putative binding sites using AutoLigand and MGLTools 1.5.7. We assigned hydrogen atoms (including polar hydrogens) and Gasteiger partial charges to each of the two aSyn-bound MMP1 poses and free MMP1. Zinc and calcium ions were removed. A grid map with 1Å spacing was then applied to the hemopexin and catalytic domains in addition to the linker region, making the docking unbiased for different binding sites, especially allosteric ones. AutoLigand was then used to define four binding site calculations per structure, beginning with 100 points to define the volume of the binding envelope, then increasing the number of points to 200, 300, and finally 400. The binding envelopes defined were analyzed using AutoDock, and results visualized using MGLTools and VMD.

### Virtual screening of small molecules against a putative site on MMP1

We performed molecular docking using AutoDock, and the virtual screening was setup with Raccoon 1.1. The receptor was considered rigid, and rotatable bonds imparted flexibility to the ligands. We used default AutoDock docking parameters (see supplementary information) to determine the smallmolecule binding affinities. A collection of 1,400 FDA-approved compounds were obtained from the ZINC15 database. We began by importing the ligand multiple structure MOL2 file into Raccoon. Each ligand was automatically converted into a separate PDBQT while settling hydrogen atoms and partial charges. We calculated a set of new grid map files with a grid spacing of 0.37Å to include the selected putative binding site from the binding site prediction. For each of the three cases (free MMP1 and two aSyn-MMP1 poses), 100 ligands with the lowest binding affinities were identified. The lead molecules’ ZINC IDs were converted to chemical IDs (CIDs) for comparison. The top 10 ligands were identified by their common names.

## Acknowledgments

One grant to S.K.S. and J.K.S. from the National Institutes of Health (RGM137295A) partially supported this work. A.N is grateful to the Oxford Science Innovation, NIHR Oxford Biomedical Research Centre, and NIHR Oxford Health Biomedical Research Centre (Informatics and Digital Health theme, grant BRC-1215-20005) for funding.

## Author contributions

S.K.S. conceived and designed the overall project, S.K. performed experiments, L.K. collected single molecule data, C.H. and D.W. performed all-atom simulations, D.W. performed Shannon entropy calculations, A.N. performed virtual screening, S.K., C.H., D.W., A.N., J.K.S., and S.K.S. analyzed data. S.K.S. wrote the manuscript. All authors edited the manuscript.

## Supplementary information

**Figure S1.**
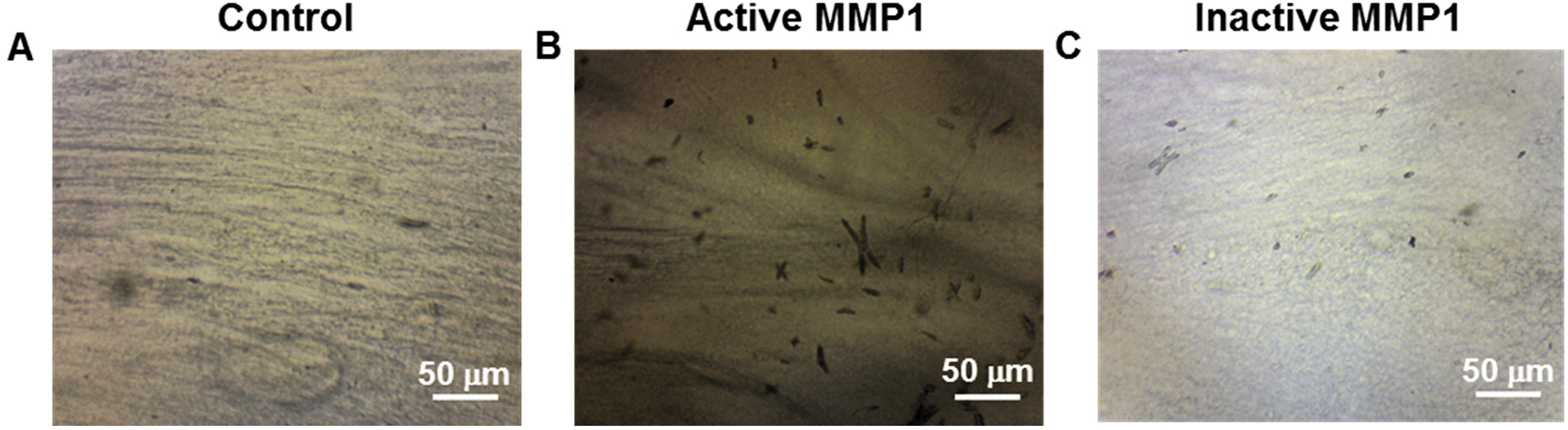
Light microscope images of aSyn-induced aggregates stained with Congo red. (**A**) Aggregates on a slide treated with protein buffer. (**B**) Aggregates on a slide treated with active MMP1 at 22 °C for 30 min. (**C**) Aggregates on a slide treated with inactive MMP1 at 22 °C for 30 min.

### Best-fit parameters for experiments

We fitted a sum of two Gaussians to the experimental histograms in **Figure 1**. The best-fit parameters are in **Table S1A**. The fit parameters b1 and b2 are the two states, S1 and S2, respectively. We fitted exponential and power-law distributions to the experimental autocorrelations in **Figure 1.** Power law distribution does not fit the experimental autocorrelations. The best-fit parameters for the exponential fits are in **Table S1B**. The kinetic rates of interconversion between S1 and S2 are in **Table S1C**. The error bars represent the standard errors of the mean.

**Table S1.** Best-fit parameters for histograms and autocorrelations in Figure 1.

**A Gaussian fit parameters for experimental histograms**
| MMP1 without ligands |  |  | MMP1 with tetracycline |  |
| --- | --- | --- | --- | --- |
|  | Active | Inactive | Active | Inactive |
| a1 | 6.03±0.05 | 1.94±0.15 | 0.64±0.04 | 1.95±0.07 |
| b1/S1 | 0.46±0.01 | 0.51±0.01 | 0.43±0.01 | 0.53±0.01 |
| c1 | 0.08±0.01 | 0.10±0.01 | 0.06±0.01 | 0.10±0.01 |
| a2 | 1.87±0.09 | 6.24±0.22 | 5.36±0.03 | 6.52±0.14 |
| b2/S2 | 0.52±0.01 | 0.58±0.01 | 0.59±0.01 | 0.61±0.01 |
| c2 | 0.04±0.01 | 0.06±0.01 | 0.10±0.01 | 0.06±0.01 |

**B Exponential fit parameters for correlations**
| MMP1 without ligands |  |  | MMP1 with tetracycline |  |
| --- | --- | --- | --- | --- |
|  | Active | Inactive | Active | Inactive |
| d | 0.20±0.01 | 0.36±0.01 | 0.53±0.01 | 0.53±0.01 |
| e | 0.08±0.01 | 0.06±0.01 | 0.05±0.01 | 0.04±0.01 |
| f | -0.01±0.01 | -0.01±0.01 | -0.01±0.01 | -0.01±0.01 |

**C Kinetic rates calculated from histograms and correlations**
| MMP1 without ligands |  |  | MMP1 with tetracycline |  |
| --- | --- | --- | --- | --- |
|  | Active | Inactive | Active | Inactive |
| k1 (s<sup>-1</sup>) | 0.0107 | 0.0396 | 0.0428 | 0.0231 |
| k2 (s<sup>-1</sup>) | 0.0708 | 0.0196 | 0.0030 | 0.0126 |

### Best-fit parameters for simulations

We considered the MMP1 dynamics as a two-state Poisson process and simulated smFRET trajectories assuming that MMP1 undergoes interconversion between two states, S1 and S2, with experimentally determined kinetic rates and noise levels. We considered active MMP1 and active site mutant of MMP1 without ligands (**Figure 1D** and **1F**). We simulated and analyzed 350 smFRET trajectories, each 1000 s long, with the input parameters in **Table S2**. The recovered parameters are on the right side of **Table S2**.

**Table S2.**
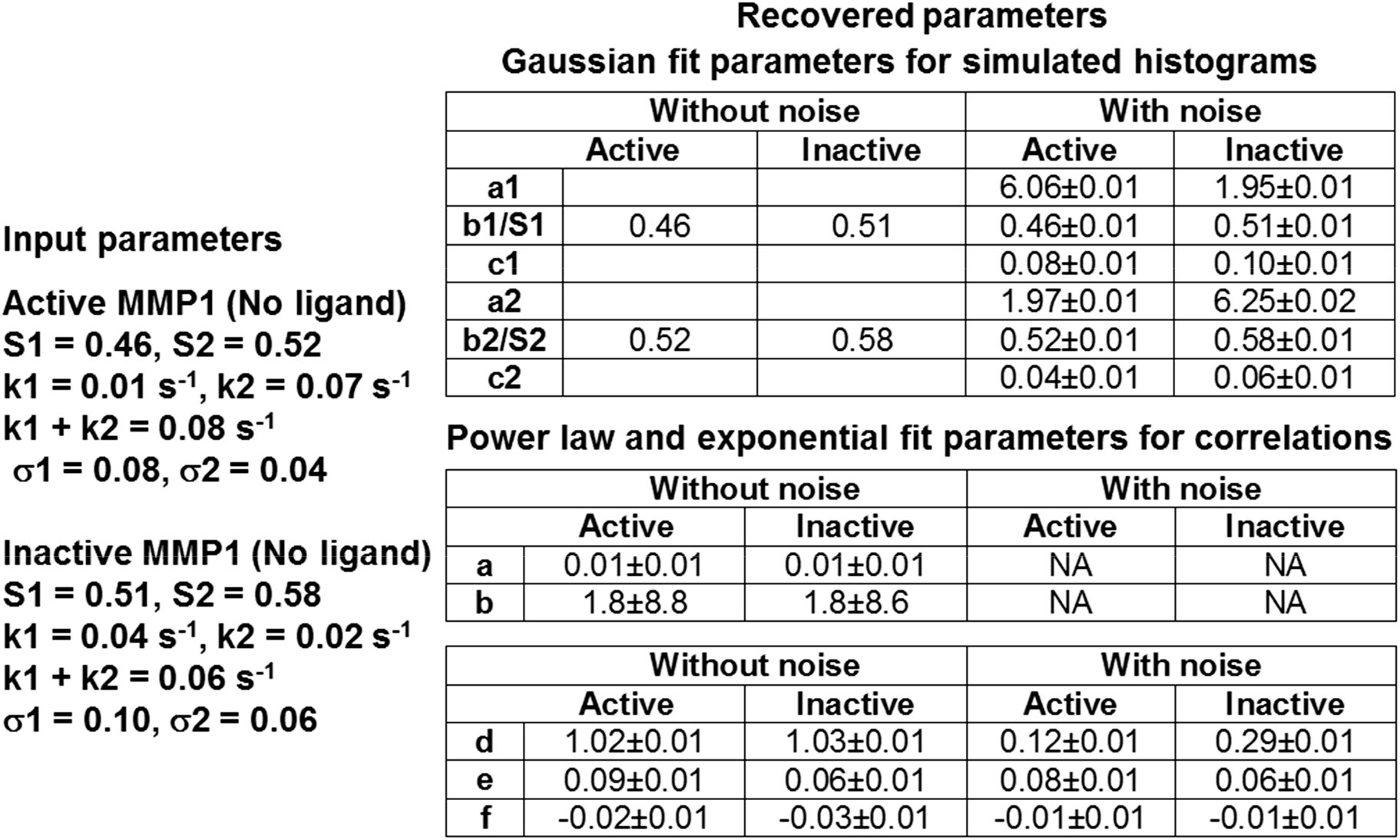
Best-fit parameters for simulated histograms and autocorrelations in Figure 2.

### RMSD stabilization of simulations

We defined the input structure at t=0 as the reference structure and calculated the root-mean-square-displacement (RMSD) to check the simulations’ stabilization. Simulations of MMP1 dynamics stabilize in ~5 ns (**Figure S2**). As such, we simulated 20 ns long dynamics for different conditions.

**Figure S2.**
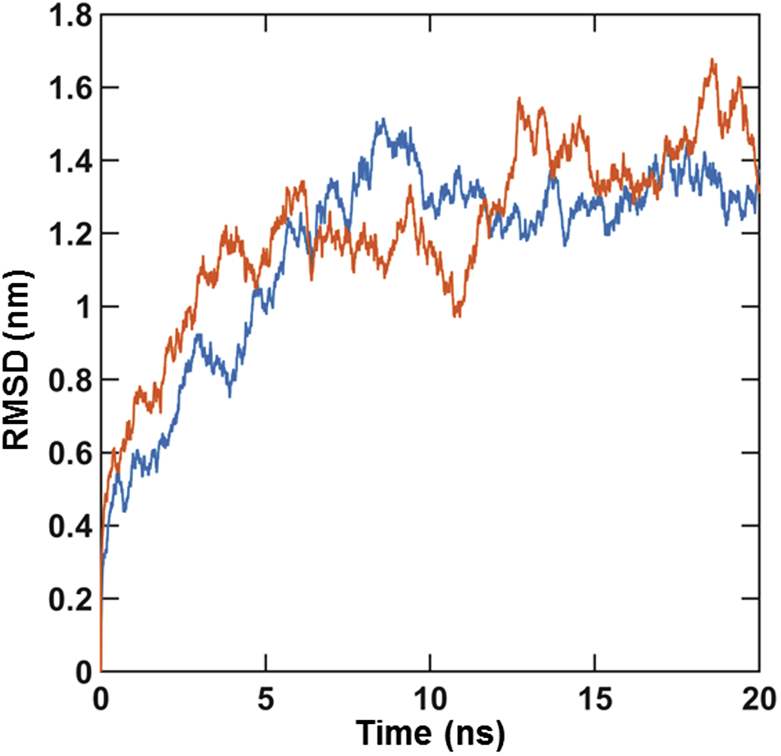
Stabilization of dynamics. RMSD for active (blue) and inactive (orange) MMP1 without ligands at 22 °C.

### Calculation of entropy

We calculated the Gray-Level Co-Occurrence Matrix (GLCM) (1) from the two-dimensional correlation plots. Then, we defined the Shannon entropy (2) from the GLCM. We describe the steps for calculating the Shannon entropy of MMP1 conformational dynamics in **Figure S3** using an arbitrary 5×5 matrix, where each element of the matrix has a value between 0 and 3. In other words, we have a 2-bit 5×5 matrix in **Figure S3**. The dimension of GLCM depends on the number of possible values (0, 1, 2, 3), i.e., 4×4 for the 5×5 matrix in **Figure S3**.

**Figure S3.**
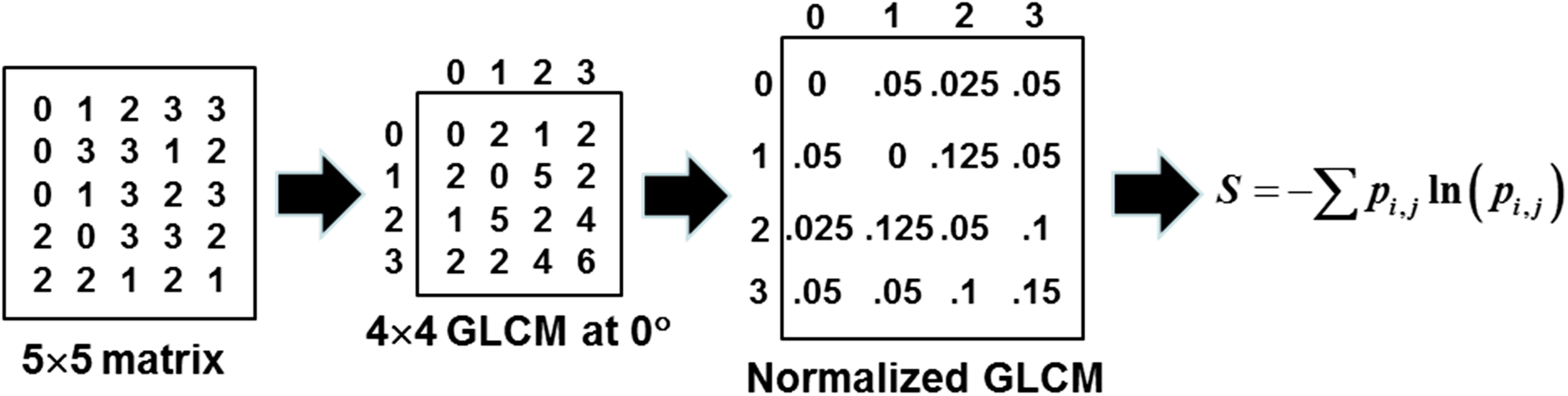
Calculation of GLCM and entropy.

We calculated the GLCM at 0°, i.e., we only considered the next neighbor on the left and right. For example, we calculated the (0,0) element of GLCM by finding the value 0s in the original 5×5 matrix and counting the number of times 0s appear on the left and right. The number is 0, and as such, the (0,0) element of GLCM is 0. To calculate the (0,1) element of GLCM, we found the value 0s in the original 5×5 matrix and counting the number of times 1s appear on the left and right. The number is 2, and as such, the (0,1) element of GLCM is 2. We repeated this process for all the elements of GLCM and obtained the matrix in the middle (Figure S3). We calculated the sum of all elements and divided each element by the sum to obtain the normalized GLCM at the left (Figure S3). We used *S* = ∑*p_i,j_* ln(*p_i,j_*), where *p_i,j_* is the (i,j) element of GLCM to quantify the Shannon entropy. By definition, we considered ln(*p_i,j_*) = 0 for *p_i,j_* = 0. Note that one can calculate GLCM at other angles (1). For MMP1 conformational dynamics, the dimension of correlation matrices is 267×267. We divided the correlation values between 0 and 1 into 10 bins of width 0.1. As such, the dimension of GLCM for MMP1 dynamics is 10×10.

**Table S3.** Best-fit parameters for simulated histograms and correlations in Figure 3.

**A Gaussian fit parameters for simulated histograms**
| Catalytic pocket opening |  |  |  |
| --- | --- | --- | --- |
|  | Pose 1 | Pose 2 | Pose 3 |
| **a1** | 1.03±0.19 | 1.44±0.03 | 3.29±2.18 |
| **b1/S1 (nm)** | 3.15±0.02 | 2.71±0.01 | 2.76±0.01 |
| **c1** | 0.12±0.03 | 0.09±0.01 | 0.11±0.01 |
| **a2** | 2.30±0.06 | 5.97±0.04 | 1.53±1.53 |
| **b2/S2(nm)** | 2.72±0.01 | 2.53±0.01 | 2.67±0.11 |
| **c2** | 0.18±0.02 | 0.07±0.01 | 0.14±0.04 |

| Interdomain distance |  |  |  |
| --- | --- | --- | --- |
|  | Pose 1 | Pose 2 | Pose 3 |
| **a1** | 0.24±0.05 | 3.65±0.04 | 3.18±1.70 |
| **b1/S1 (nm)** | 4.38±0.01 | 4.85±0.01 | 4.07±0.04 |
| **c1** | 0.05±0.01 | 0.14±0.01 | 0.13±0.02 |
| **a2** | 3.70±0.03 | 0.29±0.04 | 1.02±1.63 |
| **b2/S2(nm)** | 4.13±0.01 | 4.56±0.02 | 3.93±0.13 |
| **c2** | 0.15±0.01 | 0.11±0.03 | 0.14±0.05 |

**B Linear correlation fit parameters**
| Linear correlation |  |  |  |
| --- | --- | --- | --- |
|  | Pose 1 | Pose 2 | Pose 3 |
| **b0** | 2.72±0.13 | 3.28±0.06 | 2.72±0.05 |
| **b1** | 0.02±0.03 | -0.15±0.01 | 0.01±0.01 |

**Table S4.**
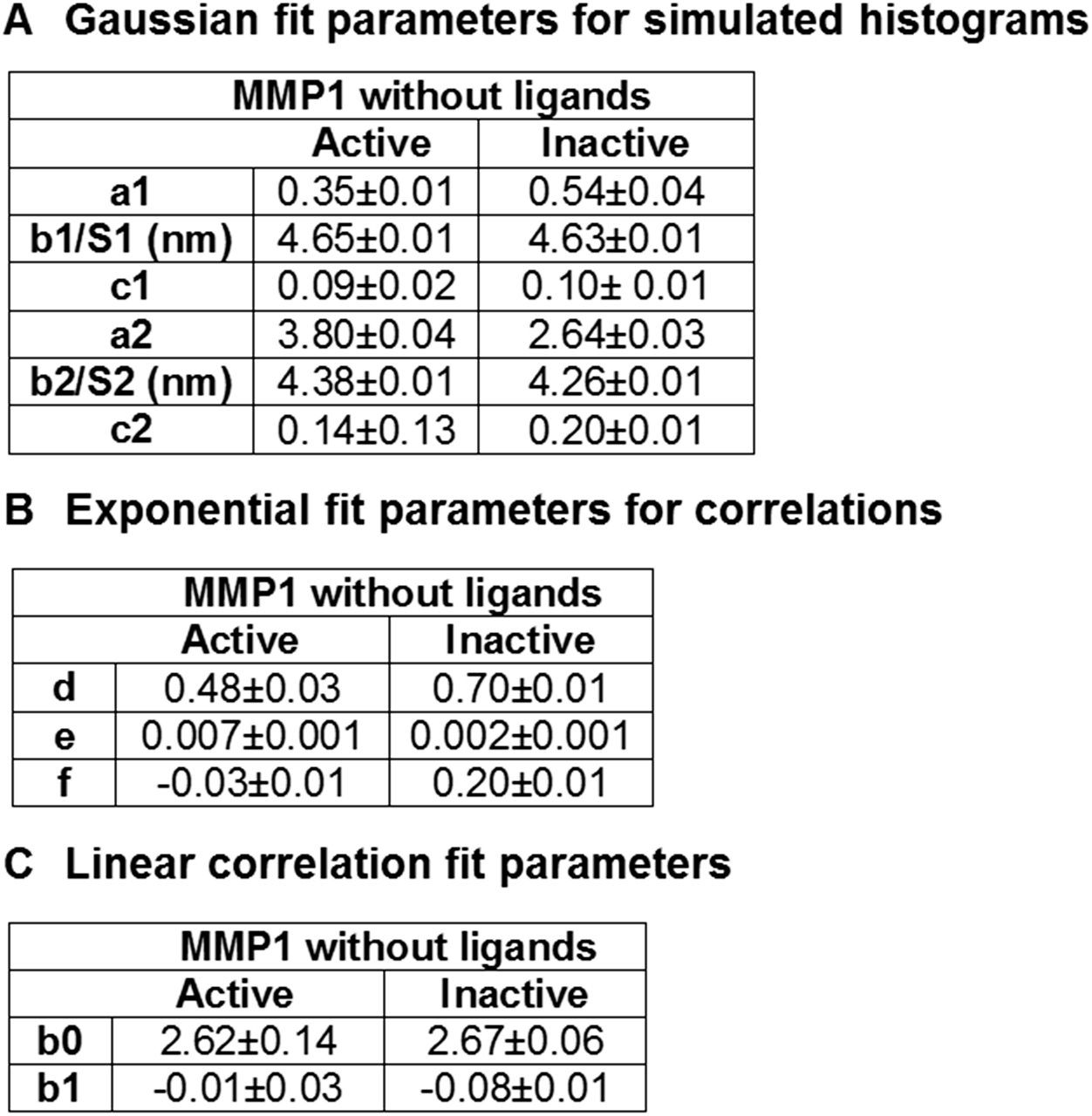
Best-fit parameters for simulated histograms and autocorrelations in Figure 5.

### Small molecule virtual screening

An example of Autodock parameters shared amongst ligands:

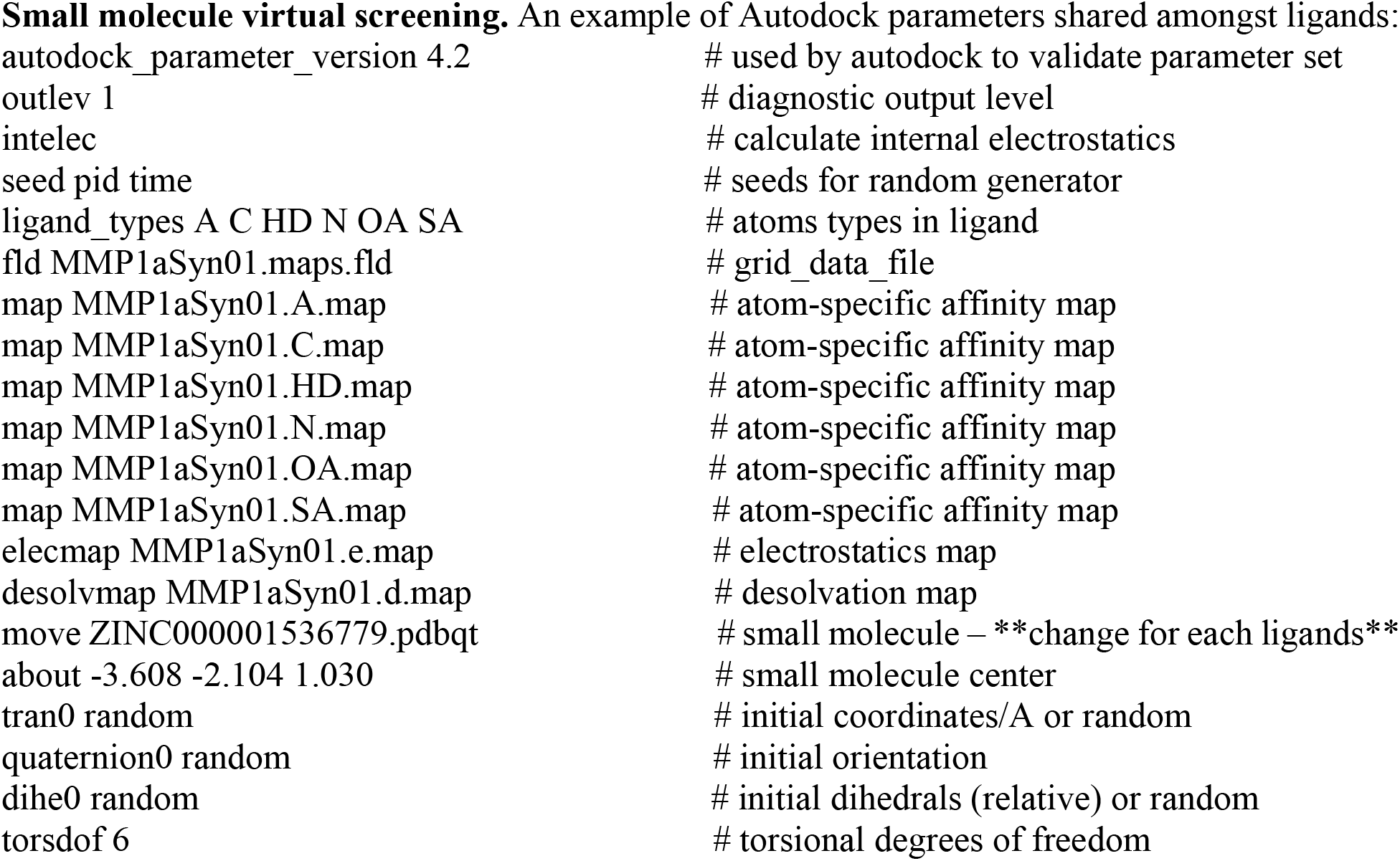
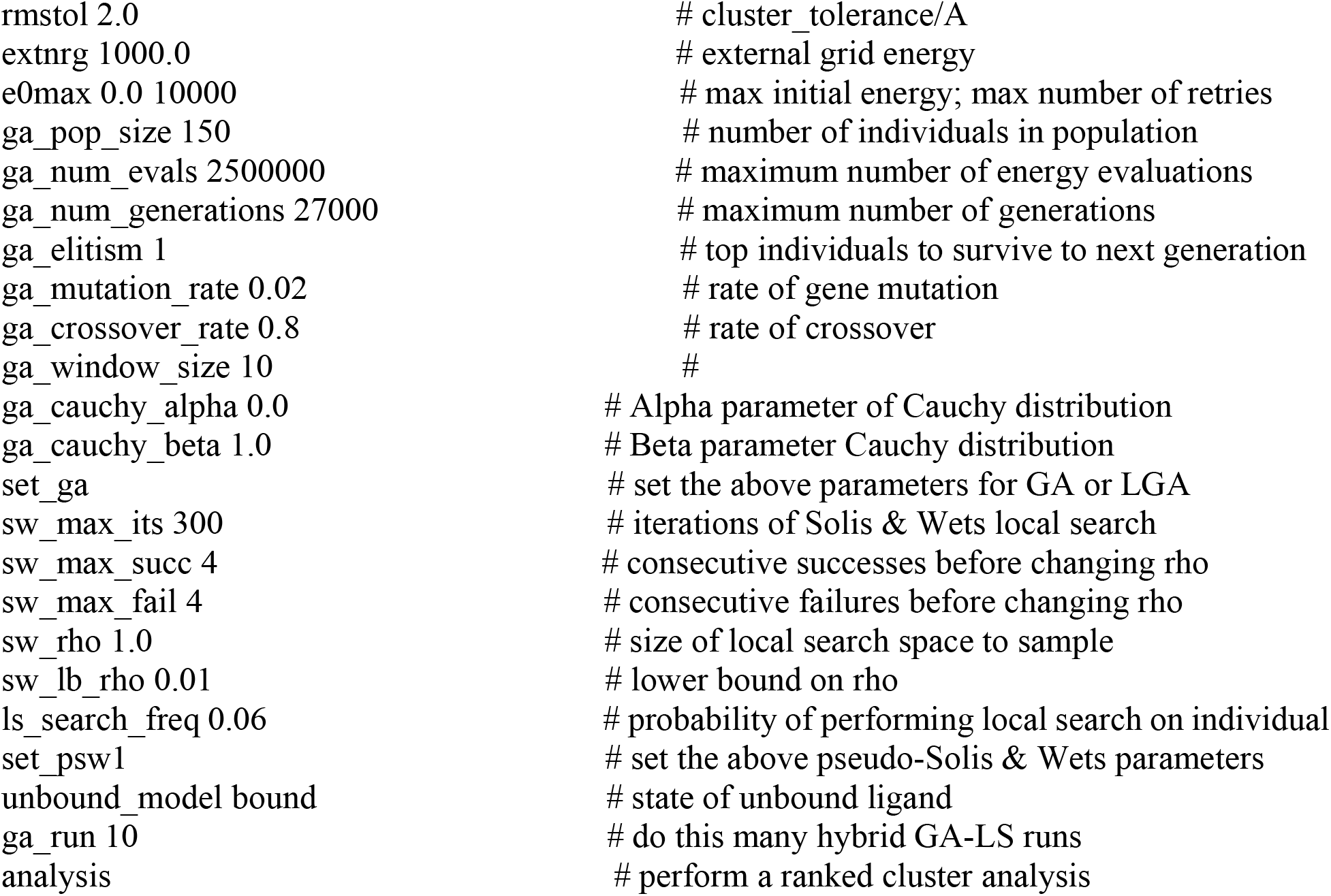

## References

[1] L. Parkkinen, T. Pirttilä, I. Alafuzoff, Applicability of current staging/categorization of α-synuclein pathology and their clinical relevance. Acta neuropathologica 115 (2008) 399–407.

[2] M.G. Spillantini, M.L. Schmidt, V.M.-Y. Lee, J.Q. Trojanowski, R. Jakes, M. Goedert, α-Synuclein in Lewy bodies. Nature 388 (1997) 839.

[3] M. Xilouri, O.R. Brekk, L. Stefanis, Alpha-synuclein and protein degradation systems: a reciprocal relationship. Molecular neurobiology 47 (2013) 537–551.

[4] I. Horvath, C.F. Weise, E.K. Andersson, E. Chorell, M. Sellstedt, C. Bengtsson, A. Olofsson, S.J. Hultgren, M. Chapman, M. Wolf-Watz, Mechanisms of protein oligomerization: inhibitor of functional amyloids templates α-synuclein fibrillation. Journal of the American Chemical Society 134 (2012) 3439–3444.

[5] D.J. Selkoe, Cell biology of protein misfolding: the examples of Alzheimer’s and Parkinson’s diseases. Nature cell biology 6 (2004) 1054.

[6] G. Forloni, L. Terreni, I. Bertani, S. Fogliarino, R. Invernizzi, A. Assini, G. Ribizzi, A. Negro, E. Calabrese, M.A. Volonté, Protein misfolding in Alzheimer’s and Parkinson’s disease: genetics and molecular mechanisms. Neurobiology of aging 23 (2002) 957–976.

[7] A.J. Espay, J.A. Vizcarra, L. Marsili, A.E. Lang, D.K. Simon, A. Merola, K.A. Josephs, A. Fasano, F. Morgante, R. Savica, Revisiting protein aggregation as pathogenic in sporadic Parkinson and Alzheimer diseases. Neurology 92 (2019) 329–337.

[8] P.T. Lansbury, H.A. Lashuel, A century-old debate on protein aggregation and neurodegeneration enters the clinic. Nature 443 (2006) 774–779.

[9] C.A. Ross, M.A. Poirier, Protein aggregation and neurodegenerative disease. Nature medicine 10 (2004) S10–S17.

[10] G.B. Irvine, O.M. El-Agnaf, G.M. Shankar, D.M. Walsh, Protein aggregation in the brain: the molecular basis for Alzheimer’s and Parkinson’s diseases. Molecular medicine 14 (2008) 451–464.

[11] H. Mizoguchi, K. Yamada, T. Nabeshima, Matrix metalloproteinases contribute to neuronal dysfunction in animal models of drug dependence, Alzheimer’s disease, and epilepsy. Biochemistry research international 2011 (2011).

[12] A. Leake, C. Morris, J. Whateley, Brain matrix metalloproteinase 1 levels are elevated in Alzheimer’s disease. Neuroscience letters 291 (2000) 201–203.

[13] J. Levin, A. Giese, K. Boetzel, L. Israel, T. Högen, G. Nübling, H. Kretzschmar, S. Lorenzl, Increased α-synuclein aggregation following limited cleavage by certain matrix metalloproteinases. Experimental neurology 215 (2009) 201–208.

[14] J.Y. Sung, S.M. Park, C.-H. Lee, J.W. Um, H.J. Lee, J. Kim, Y.J. Oh, S.-T. Lee, S.R. Paik, K.C. Chung, Proteolytic cleavage of extracellular secreted α-synuclein via matrix metalloproteinases. Journal of Biological Chemistry 280 (2005) 25216–25224.

[15] G.A. Rosenberg, Matrix metalloproteinases and their multiple roles in neurodegenerative diseases. The Lancet Neurology 8 (2009) 205–216.

[16] D. Rodríguez, C.J. Morrison, C.M. Overall, Matrix metalloproteinases: what do they not do? New substrates and biological roles identified by murine models and proteomics. Biochimica et Biophysica Acta (BBA)-Molecular Cell Research 1803 (2010) 39–54.

[17] C.J. Morrison, G.S. Butler, D. Rodríguez, C.M. Overall, Matrix metalloproteinase proteomics: substrates, targets, and therapy. Current opinion in cell biology 21 (2009) 645–653.

[18] P.G. Jobin, G.S. Butler, C.M. Overall, New intracellular activities of matrix metalloproteinases shine in the moonlight. Biochimica et Biophysica Acta (BBA)-Molecular Cell Research 1864 (2017) 2043–2055.

[19] A. Lukes, S. Mun-Bryce, M. Lukes, G.A. Rosenberg, Extracellular matrix degradation by metalloproteinases and central nervous system diseases. Molecular neurobiology 19 (1999) 267–284.

[20] B. Cauwe, G. Opdenakker, Intracellular substrate cleavage: a novel dimension in the biochemistry, biology and pathology of matrix metalloproteinases. Critical reviews in biochemistry and molecular biology 45 (2010) 351–423.

[21] S. Zucker, M. Hymowitz, C. Conner, H.M. Zarrabi, A.N. Hurewitz, L. Matrisian, D. Boyd, G. Nicolson, S. Montana, Measurement of matrix metalloproteinases and tissue inhibitors of metalloproteinases in blood and tissues: clinical and experimental applications. Annals of the New York Academy of Sciences 878 (1999) 212–227.

[22] D. Schuppan, E. Hahn, MMPs in the gut: inflammation hits the matrix. Gut 47 (2000) 12–14.

[23] J. Dzwonek, M. Rylski, L. Kaczmarek, Matrix metalloproteinases and their endogenous inhibitors in neuronal physiology of the adult brain. FEBS letters 567 (2004) 129–135.

[24] R.G. Rempe, A.M. Hartz, B. Bauer, Matrix metalloproteinases in the brain and blood–brain barrier: versatile breakers and makers. Journal of Cerebral Blood Flow & Metabolism 36 (2016) 1481–1507.

[25] R. Greenwald, L. Golub, N. Ramamurthy, M. Chowdhury, S. Moak, T. Sorsa, In vitro sensitivity of the three mammalian collagenases to tetracycline inhibition: relationship to bone and cartilage degradation. Bone 22 (1998) 33–38.

[26] T.J. Federici, The non-antibiotic properties of tetracyclines: clinical potential in ophthalmic disease. Pharmacological research 64 (2011) 614–623.

[27] M.R. Acharya, J. Venitz, W.D. Figg, A. Sparreboom, Chemically modified tetracyclines as inhibitors of matrix metalloproteinases. Drug Resistance Updates 7 (2004) 195–208.

[28] C. Balducci, G. Forloni, Doxycycline for Alzheimer’s disease: fighting β-amyloid oligomers and neuroinflammation. Frontiers in pharmacology 10 (2019).

[29] M. Bortolanza, G.C. Nascimento, S.B. Socias, D. Ploper, R.N. Chehín, R. Raisman-Vozari, E. Del-Bel, Tetracycline repurposing in neurodegeneration: focus on Parkinson’s disease. Journal of Neural Transmission 125 (2018) 1403–1415.

[30] E. Nuti, T. Tuccinardi, A. Rossello, Matrix metalloproteinase inhibitors: new challenges in the era of post broad-spectrum inhibitors. Current pharmaceutical design 13 (2007) 2087–2100.

[31] V. Mohan, D. Talmi-Frank, V. Arkadash, N. Papo, I. Sagi, Matrix metalloproteinase protein inhibitors: highlighting a new beginning for metalloproteinases in medicine. Metalloproteinases in medicine 3 (2016) 31–47.

[32] B. Sloan, N. Scheinfeld, The use and safety of doxycycline hyclate and other second-generation tetracyclines. Expert opinion on drug safety 7 (2008) 571–577.

[33] U. Eckhard, P.F. Huesgen, O. Schilling, C.L. Bellac, G.S. Butler, J.H. Cox, A. Dufour, V. Goebeler, R. Kappelhoff, U. auf dem Keller, Active site specificity profiling of the matrix metalloproteinase family: Proteomic identification of 4300 cleavage sites by nine MMPs explored with structural and synthetic peptide cleavage analyses. Matrix Biology 49 (2016) 37–60.

[34] B.I. Ratnikov, P. Cieplak, K. Gramatikoff, J. Pierce, A. Eroshkin, Y. Igarashi, M. Kazanov, Q. Sun, A. Godzik, A. Osterman, Basis for substrate recognition and distinction by matrix metalloproteinases. Proceedings of the National Academy of Sciences 111 (2014) E4148–E4155.

[35] C.M. Overall, Molecular determinants of metalloproteinase substrate specificity. Molecular biotechnology 22 (2002) 51–86.

[36] L. Kumar, A. Nash, C. Harms, J. Planas-Iglesias, D. Wright, J. Klein-Seetharaman, S.K. Sarkar, Allosteric Communications between Domains Modulate the Activity of Matrix Metalloprotease-1. Biophysical Journal 119 (2020) 360–374.

[37] L. Kumar, J. Planas-Iglesias, C. Harms, S. Kamboj, D. Wright, J. Klein-Seetharaman, S.K. Sarkar, Activity-dependent interdomain dynamics of matrix metalloprotease-1 on fibrin. Scientific Reports 10 (2020) 1–14.

[38] G. Wertheim, M. Butler, K. West, D. Buchanan, Determination of the Gaussian and Lorentzian content of experimental line shapes. Review of Scientific Instruments 45 (1974) 1369–1371.

[39] D. Kozakov, D.R. Hall, B. Xia, K.A. Porter, D. Padhorny, C. Yueh, D. Beglov, S. Vajda, The ClusPro web server for protein–protein docking. Nature protocols 12 (2017) 255.

[40] S.R. Comeau, D.W. Gatchell, S. Vajda, C.J. Camacho, ClusPro: a fully automated algorithm for protein–protein docking. Nucleic acids research 32 (2004) W96–W99.

[41] N. Yanamala, K.C. Tirupula, J. Klein-Seetharaman, Preferential binding of allosteric modulators to active and inactive conformational states of metabotropic glutamate receptors, BMC bioinformatics, BioMed Central, 2008, p. S16.

[42] N.M. Goodey, S.J. Benkovic, Allosteric regulation and catalysis emerge via a common route. Nature chemical biology 4 (2008) 474.

[43] G.G. Hammes, Multiple conformational changes in enzyme catalysis. Biochemistry 41 (2002) 8221–8228.

[44] D. Kern, E.R. Zuiderweg, The role of dynamics in allosteric regulation. Current opinion in structural biology 13 (2003) 748–757.

[45] R.D. Vale, R.A. Milligan, The way things move: looking under the hood of molecular motor proteins. Science 288 (2000) 88–95.

[46] M. Schliwa, G. Woehlke, Molecular motors. Nature 422 (2003) 759–765.

[47] S. Hammes-Schiffer, S.J. Benkovic, Relating protein motion to catalysis. Annu Rev Biochem 75 (2006) 519–541.

[48] P. Csermely, R. Palotai, R. Nussinov, Induced fit, conformational selection and independent dynamic segments: an extended view of binding events. Trends in biochemical sciences 35 (2010) 539–546.

[49] Y. Sugita, Y. Okamoto, Replica-exchange molecular dynamics method for protein folding. Chemical physics letters 314 (1999) 141–151.

[50] V.A. Jarymowycz, M.J. Stone, Fast time scale dynamics of protein backbones: NMR relaxation methods, applications, and functional consequences. Chemical reviews 106 (2006) 1624–1671.

[51] T. Schreiber, Measuring information transfer. Physical review letters 85 (2000) 461.

[52] H. Kamberaj, A. van der Vaart, Extracting the causality of correlated motions from molecular dynamics simulations. Biophysical journal 97 (2009) 1747–1755.

[53] V.M. Burger, D.J. Arenas, C.M. Stultz, A structure-free method for quantifying conformational flexibility in proteins. Scientific reports 6 (2016) 29040.

[54] T. Lenaerts, J. Ferkinghoff-Borg, F. Stricher, L. Serrano, J.W. Schymkowitz, F. Rousseau, Quantifying information transfer by protein domains: analysis of the Fyn SH2 domain structure. BMC structural biology 8 (2008) 43.

[55] Z. Wang, H. Sun, C. Shen, X. Hu, J. Gao, D. Li, D. Cao, T. Hou, Combined strategies in structure-based virtual screening. Physical Chemistry Chemical Physics 22 (2020) 3149–3159.

[56] M. Drescher, M. Huber, V. Subramaniam, Hunting the chameleon: structural conformations of the intrinsically disordered protein alpha-synuclein. ChemBioChem 13 (2012) 761–768.

[57] W.S. Davidson, A. Jonas, D.F. Clayton, J.M. George, Stabilization of α-synuclein secondary structure upon binding to synthetic membranes. Journal of Biological Chemistry 273 (1998) 9443–9449.

[58] T. Bartels, J.G. Choi, D.J. Selkoe, α-Synuclein occurs physiologically as a helically folded tetramer that resists aggregation. Nature 477 (2011) 107.

[59] E.S. Luth, T. Bartels, U. Dettmer, N.C. Kim, D.J. Selkoe, Purification of α-synuclein from human brain reveals an instability of endogenous multimers as the protein approaches purity. Biochemistry 54 (2014) 279–292.

[60] L. Kumar, W. Colomb, J. Czerski, C.R. Cox, S.K. Sarkar, Efficient protease based purification of recombinant matrix metalloprotease-1 in E. coli. Protein expression and purification 148 (2018) 59–67.

[61] S. Kamboj, C. Harms, L. Kumar, D. Creamer, C. West, J. Klein-Seetharaman, S.K. Sarkar, A method of purifying alpha-synuclein in E. coli without chromatography. Heliyon 7 (2021) e05874.

[62] X. Zhuang, L.E. Bartley, H.P. Babcock, R. Russell, T. Ha, D. Herschlag, S. Chu, A singlemolecule study of RNA catalysis and folding. Science 288 (2000) 2048–2051.

[63] J. Czerski, W. Colomb, F. Cannataro, S. Sarkar, Spectroscopic identification of individual fluorophores using photoluminescence excitation spectra. Journal of microscopy 270 (2018) 261–271.

[64] W. Colomb, J. Czerski, J. Sau, S. Sarkar, Estimation of microscope drift using fluorescent nanodiamonds as fiducial markers. Journal of microscopy 266 (2017) 298–306.

[65] A. Dittmore, J. Silver, S.K. Sarkar, B. Marmer, G.I. Goldberg, K.C. Neuman, Internal strain drives spontaneous periodic buckling in collagen and regulates remodeling. Proceedings of the National Academy of Sciences (2016) 201523228.

[66] S.K. Sarkar, B. Marmer, G. Goldberg, K.C. Neuman, Single-molecule tracking of collagenase on native type I collagen fibrils reveals degradation mechanism. Current biology: CB 22 (2012) 1047–1056.

[67] B.C. Arnold, Pareto distribution. Wiley StatsRef: Statistics Reference Online (2014) 1–10.

## References

1. R. M. Haralick, K. Shanmugam, I. H. Dinstein, Textural features for image classification. IEEE Transactions on systems, man, and cybernetics, 610–621 (1973).

2. C. E. Shannon, A mathematical theory of communication. The Bell system technical journal 27, 379–423 (1948).

